# Reproducibility and repeatability of six high-throughput 16S rDNA sequencing protocols for microbiota profiling

**DOI:** 10.1101/210880

**Authors:** Sajan C. Raju, Sonja Lagström, Pekka Ellonen, Willem M. de Vos, Johan G. Eriksson, Elisabete Weiderpass, Trine B. Rounge

**Author notes:** **Corresponding author:** Dr. Trine B. Rounge, Folkhälsan Research Center, Biomedicum 1, P.O. Box 63 (Haartmansgatan 8), 00014 University of Helsinki, Finland, E-mail address.

## Abstract

Culture-independent molecular techniques and advances in next generation sequencing (NGS) technologies make large-scale epidemiological studies on microbiota feasible. A challenge using NGS is to obtain high reproducibility and repeatability, which is mostly attained through robust amplification. We aimed to assess the reproducibility of saliva microbiota by comparing triplicate samples. The microbiota was produced with simplified in-house 16S amplicon assays taking advantage of large number of barcodes. The assays included primers with Truseq (TS-tailed) or Nextera (NX-tailed) adapters and either with dual index or dual index plus a 6-nt internal index. All amplification protocols produced consistent microbial profiles for the same samples. Although, in our study, reproducibility was highest for the TS-tailed method. Five replicates of a single sample, prepared with the TS-tailed 1-step protocol without internal index sequenced on the HiSeq platform provided high alpha-diversity and low standard deviation (mean Shannon and Inverse Simpson diversity was 3.19 ± 0.097 and 13.56 ± 1.634 respectively). Large-scale profiling of microbiota can consistently be produced by all 16S amplicon assays. The TS-tailed-1S dual index protocol is preferred since it provides repeatable profiles on the HiSeq platform and are less labour intensive.

## Introduction

Presently, there is rising interest in studying human microbiota using high throughput approaches based on 16S rRNA gene sequences. This gene is as a highly abundant, evolutionary conserved and phylogenetically informative housekeeping genetic marker (Lane et al., 1985; Tringe and Hugenholtz, 2008; Zheng et al., 2015). The composition and diversity of the human microbiota have been correlated to health and disease, although only few cases of causal relationships have been uncovered (Cho and Blaser, 2012; Human Microbiome Project Consortium, 2012; Nicholson et al., 2012; Scheithauer et al., 2016; van Nood et al., 2013).

While attention has focused on the intestinal microbiota, it is well known that the oral cavity also harbours a large microbial community that includes around 700 common bacterial species, out of which 35% are still unculturable (Dewhirst et al., 2010). Cultivation-independent molecular methods have validated these estimates, by identifying approximately 600 species or phylotypes using 16S rRNA gene sequencing techniques (Dewhirst et al., 2010; Paster et al., 2001). Oral bacteria have been linked to many oral diseases and non-oral diseases, testifying for their importance (Krishnan et al., 2017). While metagenomic studies have provided insight in the large coding capacity of the human microbiota (Li et al., 2014; Qin et al., 2010), taxonomic studies mainly rely on amplifying and analysing hypervariable regions of 16S rRNA gene sequences.

It is known that a precise assessment of the microbiota depends heavily on the hypervariable region selected, and primers used, whereas taxonomic resolution bias can arise with amplification of non-representative genomic regions (Wen et al., 2017; Zheng et al., 2015). Next generation sequencing (NGS) technology with application of barcode indexing are possible to achieve thousands of sequences from large number of samples simultaneously (Andersson et al., 2008; Hamady et al., 2008). However, reproducible identification and consistent quantification of bacterial profiles remain challenging (Ding and Schloss, 2014). Studies have shown that β-diversity metrics depicted significant correlation between oral bacterial composition for the V1–V3 and V3–V4 regions (Zheng et al., 2015). The 16S rRNA V3-V4 hypervariable region is widely used for various microbiological studies (Belstrøm et al., 2016; Fadrosh et al., 2014; Janem et al., 2017). Protocols have been developed using the dual indexing strategy to yield the greater utilization of available sequencing capacity (Kozich et al., 2013). High throughput and cost effective sequencing approaches are continuous being developed, urging researchers to use the latest technologies while abandoning the old ones. However, evaluation of new methodologies is a crucial step in conducting rigorous scientific research (Sinclair et al., 2015). This specifically applies to generating representative libraries of 16S rRNA gene amplicons.

In this study, we aimed to simplify amplification procedure and investigate barcoding efficacy with internal indices, for sequencing 16S rRNA gene amplicons relative to sequencing quality, depth, reproducibility and repeatability. Specifically, we tested high-throughput workflows for amplicon library construction of the 16S rRNA V3–V4 hypervariable region using Truseq and Nextera adapters with dual index and dual index plus 6-nt internal index. We assessed the reproducibility of the saliva microbiota for four saliva samples in triplicates using the Illumina MiSeq platform and the repeatability using nine control samples, including five replicates from a single individual, with the 1-step TS-tailed dual index protocol on the Illumina HiSeq platform.

## Materials and Methods

Saliva samples in triplicates from four volunteers were selected for this study (Fig. 1). The study was approved by the regional Ethics Committee of the Hospital District of Helsinki and Uusimaa (169/13/03/00/10). The saliva samples were collected in Oragene-DNA (OG-500) self-collection kits (DNA Genotek Inc, Canada) and mixed with stabilizing reagent within the collection tubes per manufacturer’s instructions by participants, and stored at room temperature. A protocol with an intensive lysis step using a cocktail of lysozyme and mechanical disruption of microbial cells using bead-beating was employed. Fifty ml lysozyme (10 mg/ml, Sigma-Aldrich), 6 ml mutanolysin (25 KU/ml, Sigma-Aldrich), and 3 ml lysostaphin (4000 U/ml, Sigma-Aldrich) were added to a 500 ml aliquot of cell suspension followed by incubation for 1 h at 37 °C. Subsequently, 600 mg of 0.1-mm-diameter zirconia/silica beads (BioSpec, Bartlesville, OK) were added to the lysate and the microbial cells were mechanically disrupted using Mini-BeadBeater-96 (BioSpec, Bartlesville, OK) at 2100 rpm for 1 min (Yuan et al., 2012). After lysis, total DNA was extracted using cmg-1035 saliva kit, and Chemagic MSM1 nucleic acid extraction robot (PerkinElmer).

**Fig. 1.**
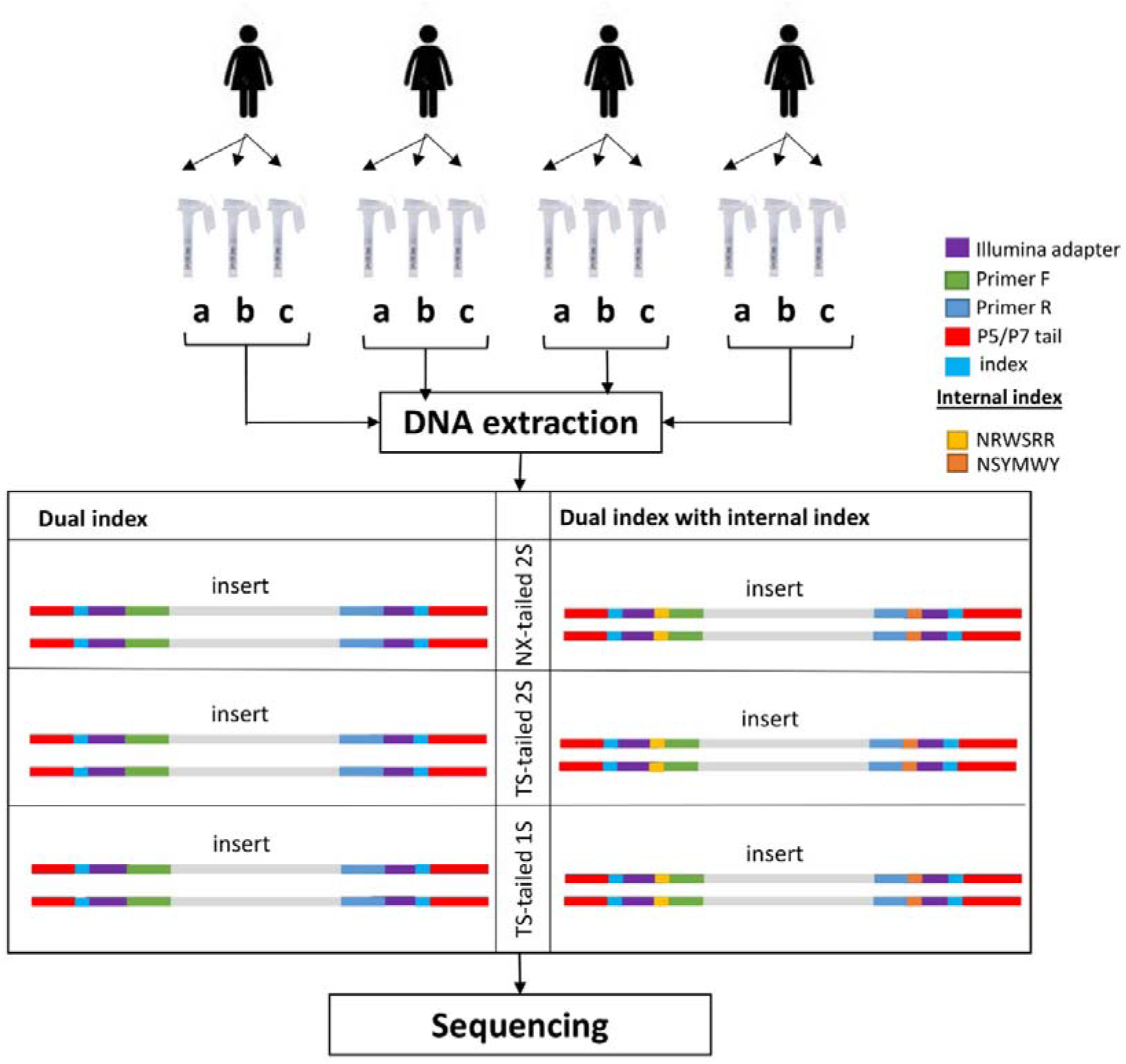
Study Design. Schematic presentation of the study design showing the number of participants and different methods implemented.

### PCR amplification

PCR amplification and sequencing libraries was prepared according to in-house 16S rRNA gene-based PCR amplification protocols. All protocols used 16S primers (S-D-Bact-0341-b-S-17: 5’ CCTACGGGNGGCWGCAG ’3 and S-D-Bact-0785-a-A-21: 5’ GACTACHVGGGTATCTAATCC 3’) targeting the V3–V4 region as reported previously (Klindworth et al., 2013). The 16S rRNA gene-based primers were modified by adding 5’ tails corresponding to the Illumina Truseq and Nextera adapter sequences to the 5’-ends. Amplification was done using primers with and without incorporated internal index (Supplementary Table S1). Two sets of index primers carrying Illumina grafting P5/P7 sequence were used: in-house index primers with Truseq adapter sequence (Supplementary Table S2) and Illumina Nextera i5/i7 adapters. All oligonucleotides (except Illumina Nextera i5/i7 adapters) were synthesized by Sigma-Aldrich (St. Louis, MO, USA).

#### TS-tailed 1-step amplification

Amplification was performed in 20 μl containing 1 μl of template DNA, 10 μl of 2x Phusion High-Fidelity PCR Master Mix (Thermo Scientific Inc., Waltham, MA, USA), 0,25 μM of each 16S primer carrying Truseq adapter, 0.5 μM of each Truseq index primer. The cycling conditions were as follows: initial denaturation at 98 °C for 30 seconds; 27 cycles at 98 °C for 10 sec, at 62 °C for 30 sec and at 72 °C for 15 sec; final extension at 72 °C for 10 min, followed by a hold at 10 °C. Separate reactions were done using 16S rRNA gene-based primers with and without incorporated internal index (denoted as ii). Here after this protocol denoted as TS-tailed-1S.

#### TS-tailed 2-step amplification

Amplification was performed in 20 μl containing 1 μl of template DNA, 10 μl of 2x Phusion High-Fidelity PCR Master Mix (Thermo Scientific Inc., Waltham, MA, USA), 0,5 μM of each 16S primer carrying Truseq adapter. The cycling conditions were as follows: initial denaturation at 98 °C for 30 sec; 27 cycles at 98 °C for 10 sec, at 62 °C for 30 sec and at 72 °C for 15 sec; final extension at 72 °C for 10 min, followed by a hold at 10 °C. Separate reactions were done using 16S rRNA gene-based primers with and without incorporated internal index. Following PCR amplification, samples were purified using a Performa V3 96-Well Short Plate (Edge BioSystems, Gaithersburg, MD, USA) and QuickStep 2 SOPE Resin (Edge BioSystems, Gaithersburg, MD, USA) according to the manufacturer’s instructions. An additional PCR step was needed to add index sequences to the PCR product. Amplification was performed in 20 μl containing 1 μl of diluted (1:100) PCR product, 10 μl of 2x Phusion High-Fidelity PCR Master Mix (Thermo Scientific Inc., Waltham, MA, USA), 0,5 μM of each Truseq index primer. The cycling conditions were as follows: initial denaturation at 98 °C for 2 min; 12 cycles at 98 °C for 20 sec, at 65 °C for 30 sec and at 72 °C for 30 sec; final extension at 72 °C for 5 min, followed by a hold at 10 °C. Here after this protocol denoted as TS-tailed-2S.

#### NX-tailed 2-step amplification

Amplification was performed in 20 μl containing 1 μl of template DNA, 10 μl of 2x Phusion High-Fidelity PCR Master Mix (Thermo Scientific Inc., Waltham, MA, USA), 0.5 μM of each of the 16S rRNA gene-based primers carrying Nextera adapters. The cycling conditions were as follows: initial denaturation at 98 °C for 30 sec; 27 cycles at 98 °C for 10 seconds, at 62 °C for 30 sec and at 72 °C for 15 sec; final extension at 72 °C for 10 min, followed by a hold at 10 °C. Separate reactions were done using 16S rRNA gene-based primers with and without incorporated internal index. Following PCR amplification, samples were purified using a Performa V3 96-Well Short Plate (Edge BioSystems, Gaithersburg, MD, USA) and QuickStep 2 SOPE Resin (Edge BioSystems, Gaithersburg, MD, USA) according to the manufacturer’s instructions. An additional PCR step was needed to add index sequences to the PCR product. Amplification was performed according to Illumina Nextera protocol to amplify tagmented DNA with following exceptions: i) reaction volume was downscaled to 20 μl, ii) 1 μl of diluted (1:100) PCR product was used as template, and iii) reaction mix was brought to the final volume with laboratory grade water. Here after this protocol denoted as NX-tailed-2S.

#### Pooling, purification and quantification

Following PCR amplifications, libraries were pooled in equal volumes. Library pool was purified twice with Agencourt® AMPure® XP (Beckman Coulter, Brea, CA, USA) according to the manufacturer’s instructions using equal volumes of the Agencourt® AMPure® XP and the library pool. The purified library pool was analyzed on Agilent 2100 Bioanalyzer using Agilent High Sensitivity DNA Kit (Agilent Technologies Inc., Santa Clara, CA, USA) to quantify amplification performance and yield.

#### Sequencing

Sequencing of PCR amplicons was performed using the Illumina MiSeq instrument (Illumina, Inc., San Diego, CA, USA). Samples were sequenced as 251x 2 bp paired-end reads and two 8-bp index reads. DNA extracted from nine blank samples, two water samples and nine control saliva samples (in which 5 samples are replicates of sample 4c) using the above mentioned protocol and amplified with TS-tailed 1S protocol without internal index, and sequencing performed (271 x 2 bp) using the Illumina HiSeq instrument.

### Phylogenetic Analysis

Sequencing quality, index trimming and length filtering was carried out using Nesoni clip Version 0.130 (https://github.com/Victorian-Bioinformatics-Consortium/nesoni). Resulting sequences were processed using MiSeq_SOP in mothur (Version v.1.35.1) (Schloss et al., 2009) and sequences were aligned to ribosomal reference database arb-SILVA Version V119 (Quast et al., 2012). We used both SILVA database and the Human Oral Microbiome Database (HOMD) database for the alignment and classification of sequences but present here only the results from the SILVA database and taxonomy as it provides comprehensive, quality checked and regularly updated databases of aligned small (16S / 18S, SSU) and large subunits (Quast et al., 2012). To obtain high quality data for analysis, sequence reads containing ambiguous bases, homopolymers > 8 bp, more than one mismatch in the primer sequence, or less than default quality score in mothur were removed. Assembled sequences with > 460 bp length and singletons were removed from the analysis. Chimeric sequences were also removed from the data set using the UCHIME algorithm within the mothur pipeline (Edgar et al., 2011). The high-quality sequence reads were aligned to the Silva 16S rRNA database (Version V119) and clustered into operational taxonomic units (OTUs) at a cut-off value > 98% sequence similarity. OTUs were classified using the Silva bacteria taxonomy reference. OTUs were calculated at distance 0.02 and alpha diversity (Shannon and inverse Simpson index) was calculated per sample. These diversity indexes are shown to be a robust estimation of microbial diversity (Haegeman et al., 2013).

#### Statistic procedures

Microbial diversity indices, both Shannon and Inverse Simpson, were used to summarize the diversity of a population. Simpson’s index is more weighted on dominant species whereas Shannon index assumes all species are represented in a sample and that they are randomly sampled (Lozupone and Knight, 2008). Kruskal-Wallis (KW-test) test was performed on the alpha diversity indices to assess the statistical significance difference between microbial diversity and the methods used. Principal coordinate analysis (PCoA) plotted with bray-curtis distance without normalizing the data using biom formatted OTUs from mothur to the phyloseq R-package Ver 1.22.3 (McMurdie and Holmes, 2014). Intraclass correlation coefficients (ICC) to quantify the reproducibility, stability, and accuracy or neutrality of different protocol for six metrics included relative abundances of four major phyla (Actinobacteria, Bacteroidetes, Firmicutes and Proteobacteria) and two alpha diversity indices (Shannon & Inverse Simpson index). The ICCs were estimated using the SPSS (version 22) based on the mixed effects model (Sinha et al., 2016). All the graphics and plots were made in R using ggplot2 package (Wilkinson, 2011).

## Results

### 16S rRNA sequencing

Saliva microbiota sequence data of the 16S rRNA V3-V4 region for 4 individuals in triplicates using TS-tailed and NX-tailed amplification, with and without internal index, were collected on the Illumina MiSeq platform (Table 1).

**Table 1:** Sequencing statistics, quality check passed sequences and sequences classified for samples (combined for methods used) sequenced in MiSeq and HiSeq platform.

| #samples | Protocol | Total sequences | Reads/sample<br>(Min) (Max) (Mean) |  |  | QC passed sequences | #Sequences classified | % of sequences |
| --- | --- | --- | --- | --- | --- | --- | --- | --- |
| 12 | NX-tailed-2S | 56032 | 2273 | 6480 | 4669 | 52070 | 34332 | 61.27 |
| 12 | NX-tailed-2S ii* | 64791 | 2452 | 10192 | 5399 | 55407 | 24836 | 38.33 |
| 12 | TS-tailed-2S | 115736 | 6940 | 15161 | 9644 | 100788 | 69783 | 60.29 |
| 12 | TS-tailed-2S ii | 142880 | 8048 | 16271 | 11906 | 121134 | 66014 | 46.20 |
| 12 | TS-tailed-1S | 96252 | 733 | 10868 | 8021 | 79339 | 56761 | 58.97 |
| 12 | TS-tailed-1S ii | 123165 | 2187 | 14519 | 10263 | 89631 | 48479 | 39.36 |
| 9 | TS-tailed-1S ** | 1228989 | 48100 | 398420 | 136554 | 974702 | 711088 | 57.86 |

\* Internal indices, \*\* Control samples sequenced on the HiSeq platform

Two control water samples, nine saliva control samples (including 5 replicates) and blank samples using TS-tailed 1S protocol without internal index, were sequenced on the Illumina HiSeq platform. Samples sequenced using TS-tailed 1S and 2S protocol with and without internal index generated comparatively higher amounts of sequence reads. This was true also after trimming of low-quality sequences (Fig. 2 and Supplementary table S3). The sequences were clustered and assigned to 1086 OTUs. Sequence coverage and percentage of sequences passed quality check from each protocol and qualified for taxonomic classification are summarized in Table 1.

**Fig. 2:**
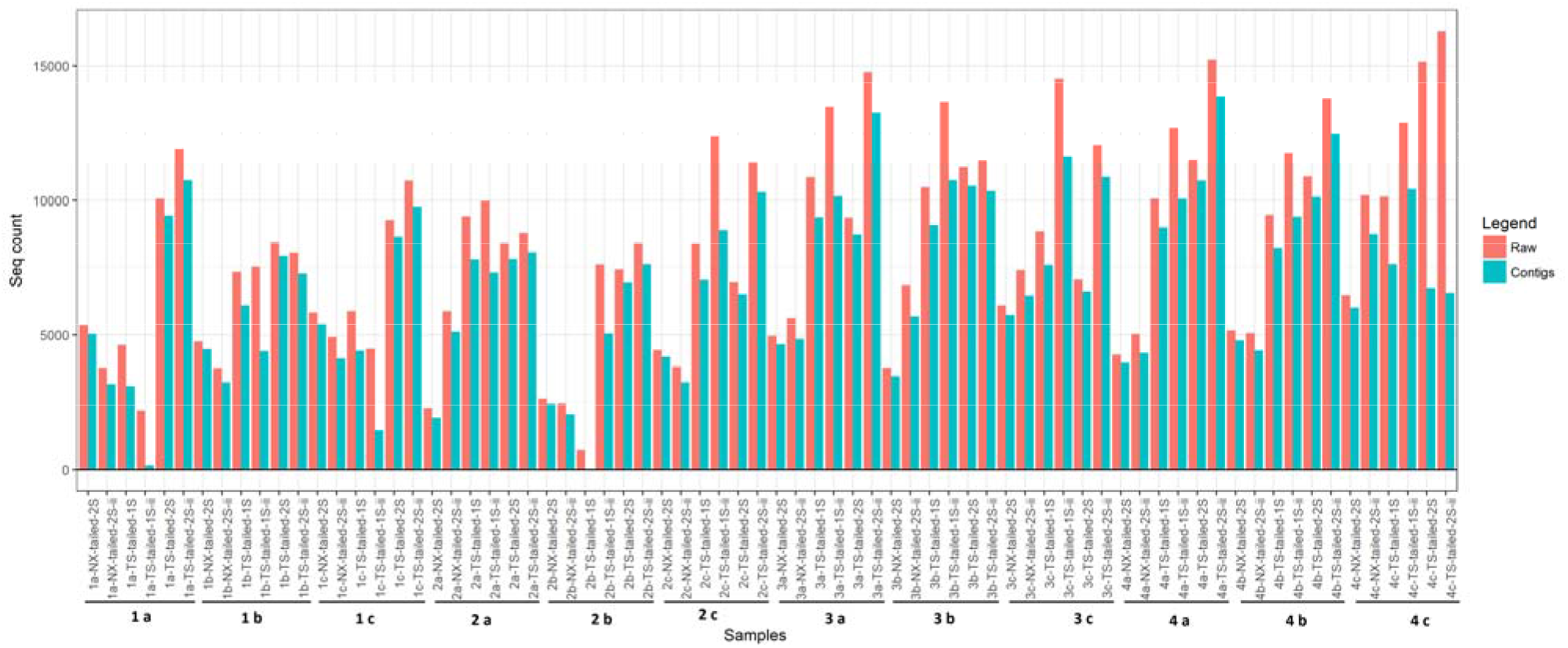
Sequence data filtering. Distribution of sequences before (Raw data count in blue colour) and after (contigs count in red colour) quality check and assembly for the saliva microbiota sequenced in MiSeq platform.

The protocols with the internal index approach showed consistently 14-23 % lower OTU’s per sample for all protocol. About 61% of saliva microbiota sequences remained after quality check using NX-tailed protocol without internal index, while only 38% remained using NX-tailed protocol with internal index. In our study, the NX-tailed protocol produced slightly less sequences than the TS-tailed protocols, with 4669 and 5399 mean reads per sample respectively. About 60% of saliva microbiota passed the quality check in TS-tailed without internal index and produced in our protocol more than 8000 reads per sample. Principle coordinate analysis (PCoA) using Bray-Curtis distance, to visualize broad trends of how similar or different bacteria are between triplicate samples, shows that samples cluster according to the individual (Fig. 3).

**Fig. 3:**
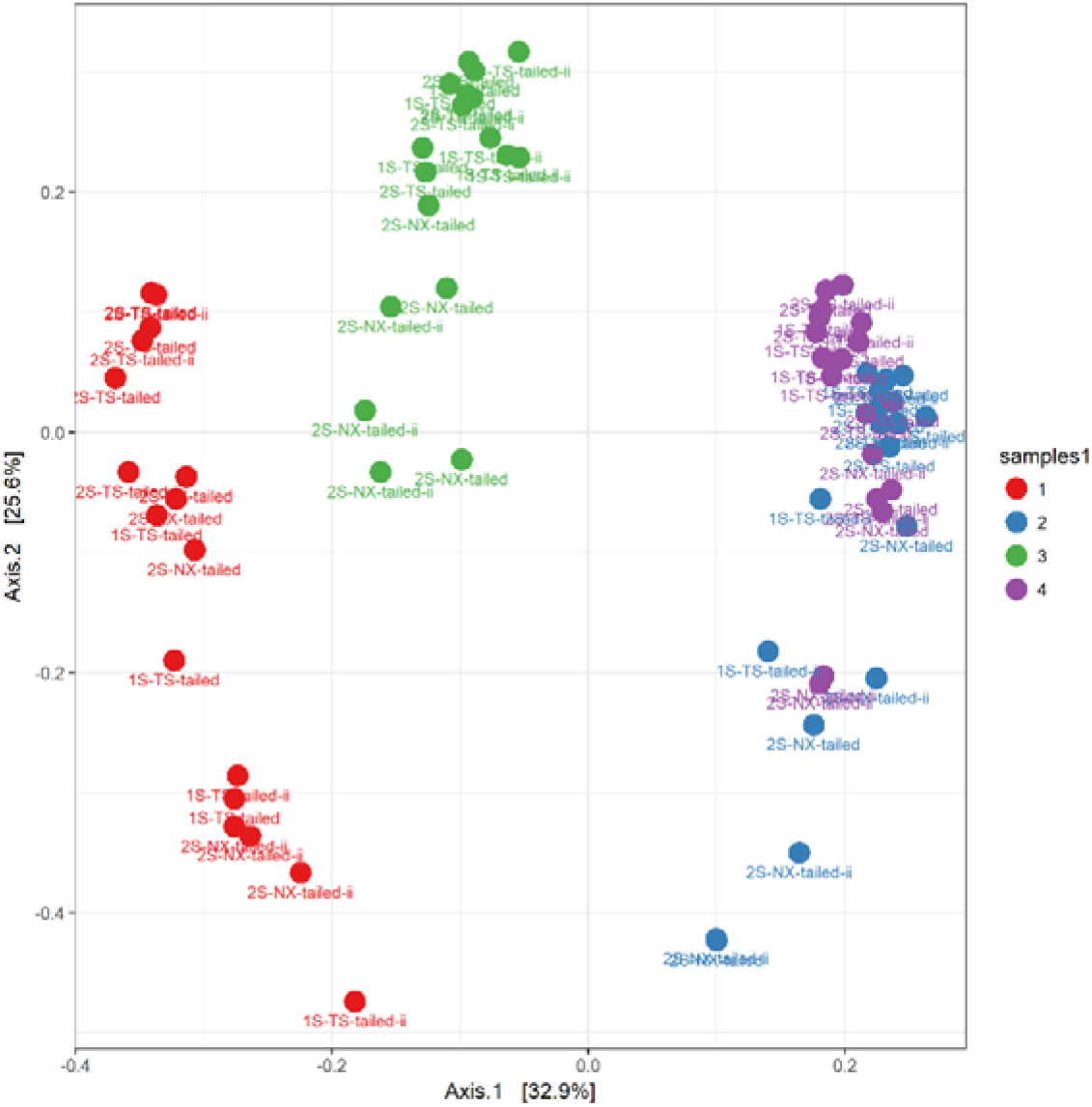
Principal coordinate analysis (PCoA) plot. PCoA plot, based on the bray-curtis distances. The percentage of the total variance explained by each PC is indicated in parentheses.

### Alpha diversity of saliva microbiota is similar for all the protocols

The Shannon diversity and inverse Simpson indices used to calculate the alpha diversity showed similar diversity for each sample irrespective of the protocols used with exception of few outliers (Fig. 4 a and b). The outliers are the samples with low diversity and low sequence depth, < 4000 sequences. Though Shannon diversity index showed less variation according to the sequence depth compared to inverse-Simpson index, we did not find any significant relationship (KW-test) between the diversity indices and the protocols used.

**Fig. 4 a & b:**
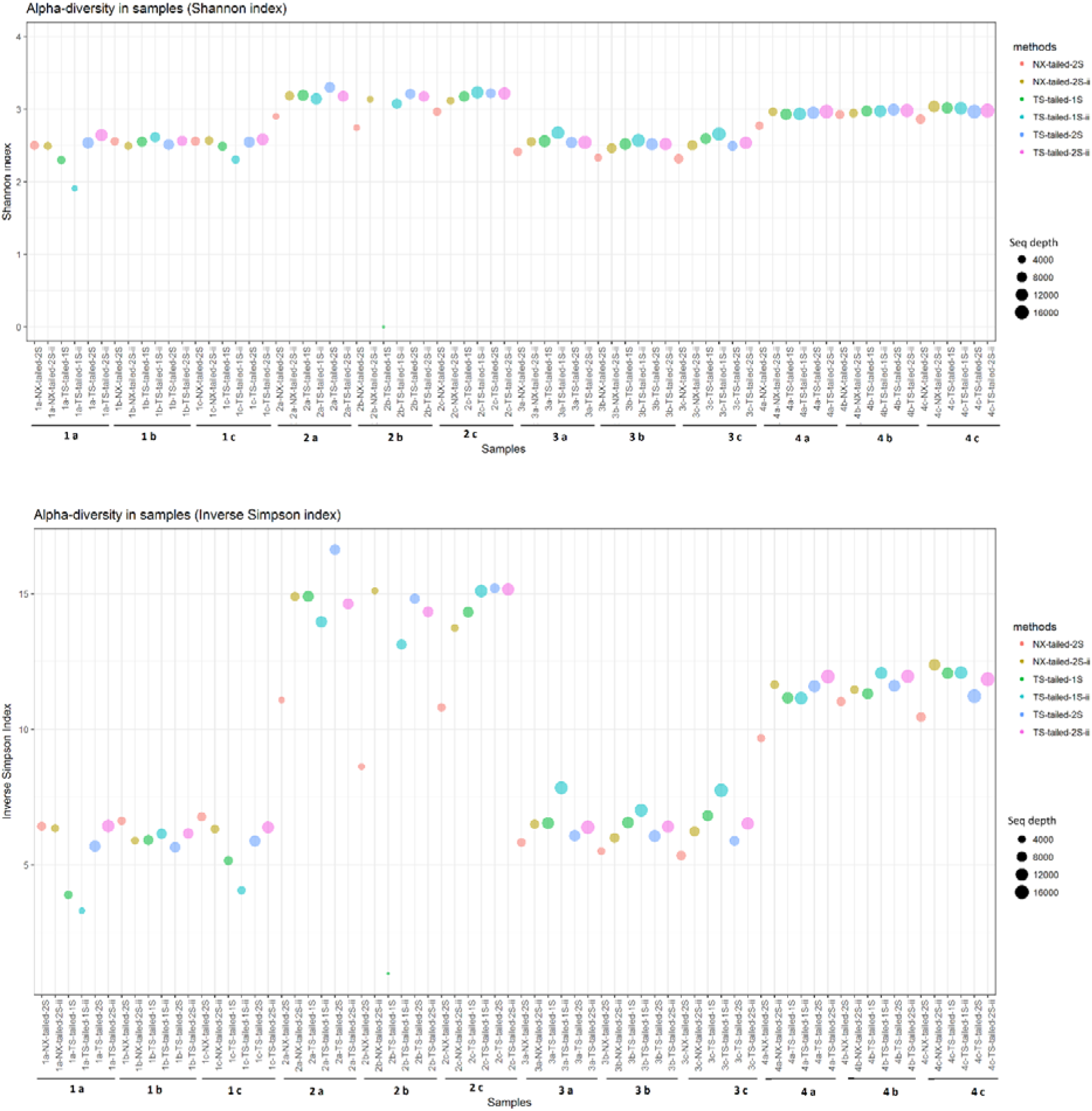
Microbial diversity in saliva. Alpha diversity measured using Shannon (a) and Inverse Simpson (b) index in each replicates. Bubble size depicts the sequence depth and bubble colour is Illumina method used.

### Consistent occurrence of bacterial abundance within the protocols

Taxonomic composition of saliva microbiota from four samples with different amplification protocols with and without internal index showed sample specific composition profile at two taxonomic levels. The bacterial relative abundance at phylum level was measured using the top five abundant phyla; Actinobacteria, Bacteroidetes, Firmicutes, Fusobacteria and Proteobacteria (Fig. 5).

**Fig. 5:**
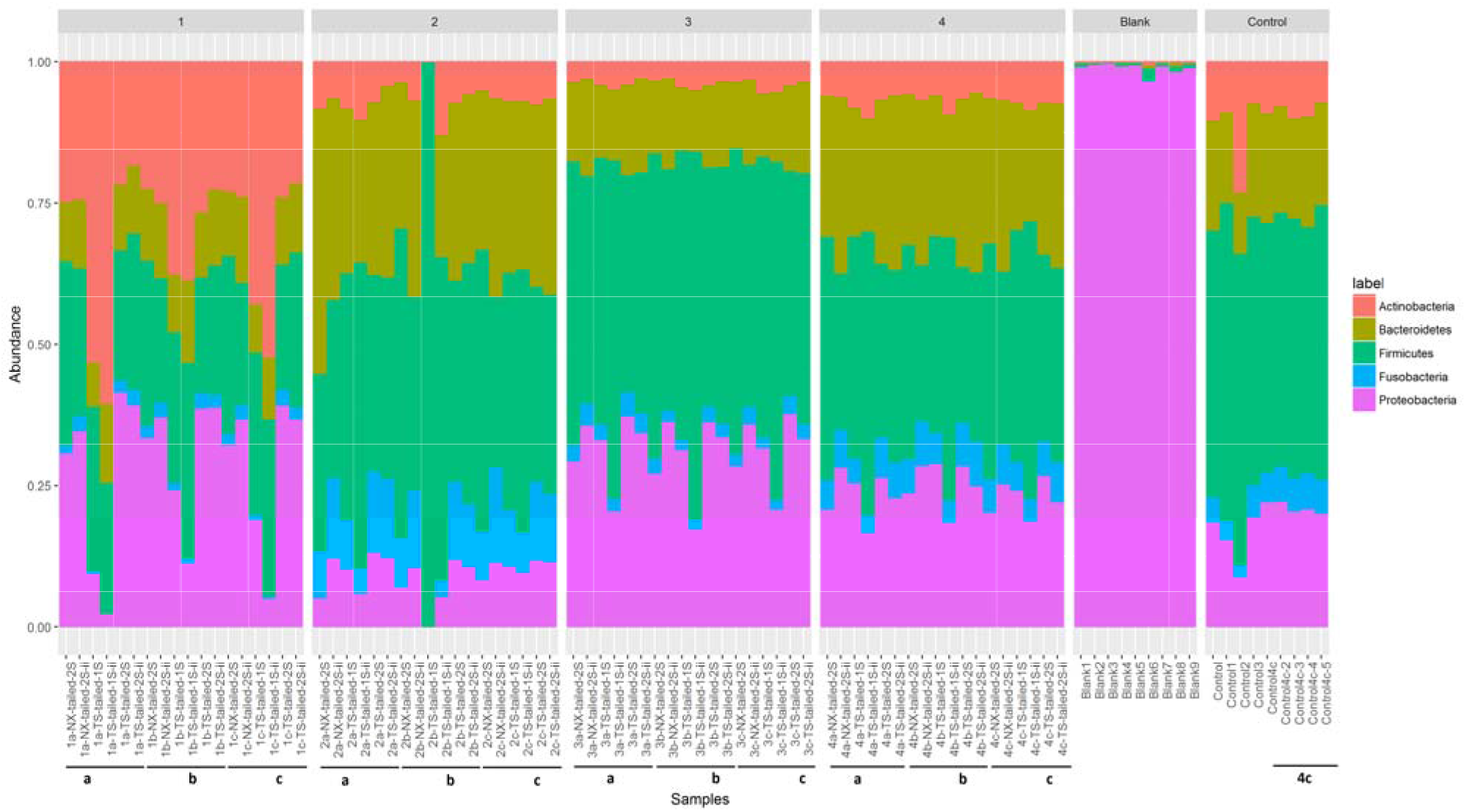
Phylum abundance. Composition of abundant phylum in each sample separated by individuals. Phylum composition of blank and control samples were also included in the figure as separate samples. Samples with low sequence depth are marked in red coloured box.

Similar patterns of phyla abundance were observed for the samples from same individuals using the different protocols. However, detailed comparison of the phyla abundance showed that the oral microbiota of individual 1 included a high abundance of Actinobacteria, Proteobacteria and Firmicutes, that of individual 2 included mainly Firmicutes and Bacteroidetes, whereas that of individuals 3 and 4 included mainly Firmicutes, Bacteroidetes and Proteobacteria. Sample 2b from individual 2 which was sequenced using the TS-tailed 1S protocol without internal index was an outlier with only 733 sequences. The relative abundance at the bacterial genus level was measured using the top 30 abundant genera (Supplementary Figure. S4). Similar patterns of genus abundance were also observed for the samples from same individuals using the different protocols. However, these compositions differed between the individuals in line with the differences at the phylum level (Fig. 5).

### Reproducibility and stability of the protocols

Average Shannon diversity for sample 1 was comparatively similar except for the TS-tailed 1S protocol with internal index. In sample 2, NX-tailed 2S and TS-tailed 1S without internal indices protocol yielded comparatively less Shannon diversity. Where as in sample 3 and sample 4 Shannon diversity was comparatively similar for all the protocols. Average Inverse Simpson diversity was comparatively less, using NX-tailed 2S protocol for sample 2, 3 and 4, TS-tailed 1S protocol for sample 1, TS-tailed 1S protocol in sample 1 and 2, and, TS-tailed 1S protocol with internal index for sample 1 (Supplementary Table S5). Intra-class correlation coefficients (ICC) used to enumerate the reproducibility and stability of different protocols for six metrics included relative abundances of four top abundant phyla and two alpha diversity indices showed comparatively better reproducibility and stability with TS-tailed 2S protocol with and without internal index (Fig. 6).

**Fig. 6:**
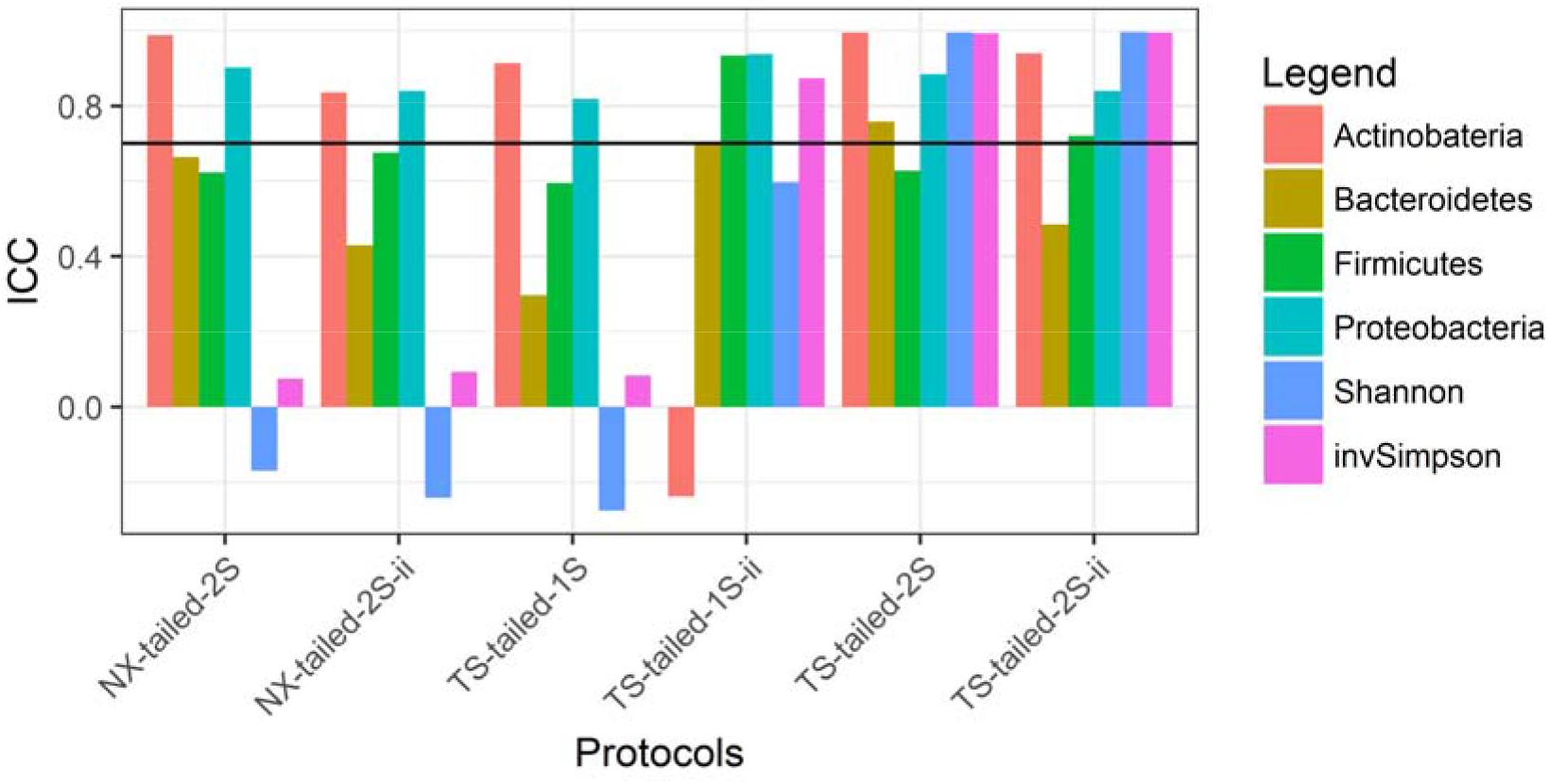
Reproducibility and stability of the protocols. Intra-class correlation coefficient plotted for six metrics included relative abundances of four major phyla (Actinobacteria, Bacteroidetes, Firmicutes and Proteobacteria) and two alpha diversity indices (Shannon & Inverse Simpson index).

Actinobacteria from TS-tailed 1S protocol with internal indices, and Shannon index from TS-tailed 1S, NX-tailed protocol, and NX-tailed protocol with internal index showed negative ICC.

### Repeatability of the saliva microbiota with TS-tailed 1S protocol

Repeatability of the saliva microbiota using the TS-tailed 1S protocol, which give the reproducibility and stability, were tested with nine control samples in HiSeq Illumina platform. We also amplified and sequenced negative controls; nine blank samples and two water samples to identify reagents and laboratory contamination. HiSeq platform provided 28936 mean sequences data for nine blank samples, 136554 sequences for nine control samples and 790 mean sequences for water samples. The result showed low diversity for blank samples sequenced and high diversity for the control samples (Fig. 7). None of the sequences from the water samples could be assembled. Mean Shannon diversity was 0.374 and 3.15 and standard deviation (SD) of 0.122 and 0.097, for blank and control samples respectively. Whereas mean inverse Simpson diversity was 1.177 and 13.460 and SD of 0.097 and 1.634, for blank and control samples respectively. Two abundant OTUs from the blank samples were explicitly assigned to two genera of the Proteobacteria phylum, *Pseudomonas* and *Achromobacter*. Bacterial relative abundance of control samples at phyla level shows high abundant of phyla Firmicutes, Bacteroidetes and Proteobacteria (Fig. 5). Relative abundance of bacteria at genus level showed that the control samples were enriched in *Veillonella, Prevotella, Rothia, Neisseria* and *Fusobacterium* spp. (Supplementary Figure. S4).

**Fig. 7 a & b:**
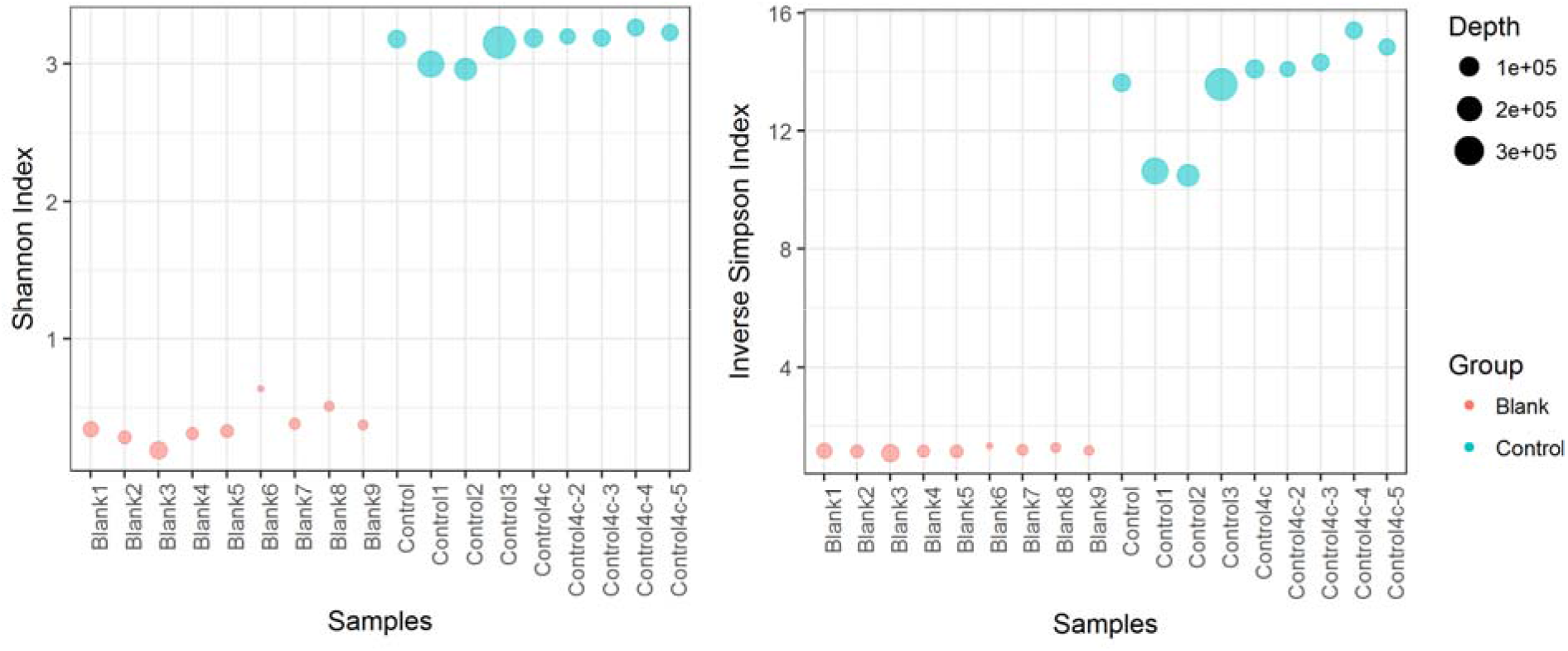
Microbial diversity in blank and control samples. Alpha diversity measured using Shannon (7a) and Inverse Simpson (7b) index in blank and control samples data from HiSeq platform.

## Discussion

Several studies have successfully used the Illumina technology approaches for 16S rRNA gene amplicon sequencing on diverse sample types (Bartram et al., 2011; Caporaso et al., 2012; Claesson et al., 2010; Degnan and Ochman, 2012; Fadrosh et al., 2014; Gloor et al., 2010; Kozich et al., 2013; Sinclair et al., 2015; Zhou et al., 2011). However, protocols differ in extraction methods, primers, chemistry and sequencing length between studies and a gold standard has not been established. In this study, we compared the reproducibility of six Illumina technology based amplification protocols on saliva samples with primers that were modified in-house (Klindworth et al., 2013; Yuan et al., 2012). We aimed to simplify amplification procedure, investigate barcoding efficacy and expand the number of available barcodes to make 16S assays feasible to run large smaple sets on the HiSeq platform.

Cells may vary in their susceptibility to lysing methods. Various studies have shown that mechanical lysis gives highest bacterial diversity in 16S rRNA gene based studies, notably when communities carry hard to lyse Gram-positive bacteria, such as in faecal samples (Robinson et al., 2016; Salonen et al., 2010; Santiago et al., 2014; Yuan et al., 2012). However, oral samples extracted using either mechanical or enzymatic lysis steps showed an overall similar microbiota profiles (Lazarevic et al., 2013). A recent study also showed that saliva sample collection, storage and genomic DNA preparation with enzymatic-mechanical lysis does not significantly influence the salivary microbiome profiles (Lim et al., 2017). All samples in this study were lysed with an identical protocol including both enzymatic and mechanical disruption of microbial cells using bead-beating to reduce the bias may arise due to the lysis step.

The four saliva samples in triplicates analysed in MiSeq using the different protocols provided comparatively high sequencing coverage for the TS-tailed protocols (>10 k) and less for all other protocols (< 10 k). With current read length of 251 x 2 bp, the V3-V4 region of the rRNA gene is a possible target for sequencing (Mizrahi-Man et al., 2013), although satisfactory quality of the overlap of the forward and reverse paired-end reads may be challenging. However, extending the sequencing cycle up to 271 bp on the Illumina HiSeq platform provided ample overlap, and assembling these reads increases the reliability and quality in the overlapping region. In MiSeq, overall, 39% - 68% of the reads were discarded due to the low-quality score, unassembled pairs, assembled pairs with mismatched barcodes, minimum overlap length and archaeal or eukaryotic sequences. Quality trimming of the NX-tailed protocol sequence data discarded a lower number (6% -8%) of data though it yielded fewer sequences than the TS-tailed protocols. Protocols without internal index pairs gave comparatively high percent (>59 %) qualified for the OTU classification per sequence. Saliva samples amplicons processed with internal index pairs had lower OTU classification per sequence. Several studies have shown that high incidence of mismatching barcodes is a main loss factor in the microbiota sequencing studies (Degnan and Ochman, 2012; Sinclair et al., 2015). This suggests that the fragment length is at the borderline of what will yield high quality sequence for the overlap between the read pairs and adding only a few extra base pairs to the fragment will reduce output quality. Whereas protocols with and without internal index pairs produced different sequence depth and quality data, all the protocols provided similar bacterial profiles for each samples. Studies have reported that, with the dual-index approach large number of samples can be sequenced using a number of primers equal to only twice the square root of the number of samples (Kozich et al., 2013). Dual indexing protocol is modified by adding the heterogeneity spacers to increase nucleotide diversity at the start of sequencing reads (Fadrosh et al., 2014). Dual indexing strategy further modified by adding third Illumina compatible index with variable length heterogeneity spacers to minimizes the need for PhiX spike-in (de Muinck et al., 2017). However, the advantage of dual index with internal index is to reduce the PCR amplification artefacts in high multiplex amplicon sequencing (Peng et al., 2015) and to reduce the cost of sequencing when the study includes a large sample size.

Our results show that low amounts of sequences usually correlate with low diversity. Our sample size was not large enough to conclude that low amounts of sequences was due to the quality or quantity of DNA, technical issues in the lab or difference in robustness of the methods. However, differences in yields using the same DNA, for example seen in sample 1a and 2b, suggest that protocol robustness may cause differences in sequencing yield (Supplementary Figure S6). Laboratory protocol, sequencing platform or error rate and bioinformatics approach can be reason for the majority of variability detected in microbiota studies (Salter et al., 2014; Sinha et al., 2015) but in our study all the protocols delivered overall similar profile of the microbes in the given saliva samples in triplicates. Three OTUs were explicitly assigned only to blank samples in HiSeq run. Negative control samples often yield contaminating bacterial species which may be due to contamination of bacterial DNA in the kits used (Salter et al., 2014). This study also reported that the presence of contaminating sequences is dependent on the amount of biomass in the samples; however, we could not assess this in our samples.

Technical challenges have been reported in 16S rRNA amplicon sequencing, such as biases in estimation of population abundance in microbial communities due to the PCR primer selection, PCR template concentration and amplification conditions, pooling of multiple barcodes and sequencing. Hence, it is important to carefully interpret the experimental results from the technical replicates to validate the reproducibility of the methods (Wen et al., 2017). Average alpha-diversity indices for each samples in different protocols yielded comparatively similar profiles with one or two exception, which may due to the low sequence depth. We used the mixed-effect model–based ICC to quantify the reproducibility and stability of the Illumina MiSeq sequencing of saliva microbiome. ICC measures the variability among the multiple measurements for the same sample and assumes that the errors from different measurements have exactly the same statistical distributions and are indistinguishable from each other (Sinha et al., 2016). In our study, based on ICC, sequencing protocols using TS-tailed 2S protocol with and without internal index performed better than NX-tailed protocol and TS-tailed 1S protocols. The negative ICC values observed for all the NX-tailed protocol and TS-tailed 1S protocols may be due to high variation within a subject.

Saliva samples sequenced on HiSeq platform yielded high sequence depth ie; 48k – 398k sequences. Variation in technical replicates and low reproducibility, can be overcome by increasing the sequencing depth (Wen et al., 2017), obtainable by the HiSeq platform. Repeatability of the TS-tailed 1S method without internal index for nine control samples sequenced in HiSeq platform was given comparatively high alpha diversity and low variation (SD) among the samples. Alpha diversity was similar for the sample 4 sequencing repeated in MiSeq and HiSeq platform which support the repeatability of method TS-tailed without internal index as good protocol for microbiome studies. The major limitation of this study is the small number of samples tested for each method. However, we believe that, the number of samples and the depth of the sequencing is sufficient to identify method that should not be used, and also indicate the preferred method to use in large scale studies.

In conclusion, NX-tailed 2S protocol and TS-tailed both 1S and 2S protocols were able to reproduce bacterial profiles for the samples sequenced, however, in our hands the reproducibility was comparatively highest for the TS-tailed 2S protocols without internal index on the MiSeq platform. Repeatability of the TS-tailed 1S protocol without internal dual index for nine control samples provided high alpha diversity and little variation among the samples. Considering the cost and time efficiency of using this simplified protocol with numerous barcodes suitable for the HiSeq platform, we suggest that the TS-tailed 1S method can be considered the most effective protocol for consistent quantification of bacterial profiles in saliva. Reproducibility and repeatability should be taken into consideration in design of a large-scale epidemiological study using saliva microbiota.

## Acknowledgements

We thank the individuals who participated in this study, and the FIMM biobank and FIMM tech centre. We also thank Timo Miettinen from FIMM tech centre for helping with the internal index setup. We also thank our group members for assisting with the fieldwork of the study Nina Jokinen, Jannina Viljakainen, Stephanie von Kraemer, and the scientific advisors’ Dr Eva Roos and Professor Anna Elina Lehesjoki.

## Ethics approval and consent to participate

The study was approved by the regional Ethics Committee of the Hospital District of Helsinki and Uusimaa (169/13/03/00/10).

## Availability of data and materials

The datasets generated during the current study are available in the NCBI-SRA repository, with the accession number SRP117317.

